# Mean amplitude of low frequency fluctuations measured by fMRI at 11.7T in the aging brain of mouse lemur primate

**DOI:** 10.1101/2022.12.21.521367

**Authors:** Clément M. Garin, Marc Dhenain

## Abstract

Non-human primates are a critical species for the identification of key biological mechanisms in normal and pathological aging. One of these primates, the mouse lemur, has been widely studied as a model of cerebral aging or Alzheimer’s disease. The amplitude of low-frequency fluctuations of blood oxygenation level-dependent (BOLD) can be measured with functional MRI. Within specific frequency bands (*e.g*. the 0.01–0.1 Hz), these amplitudes were proposed to indirectly reflect neuronal activity as well as glucose metabolism. Here, we first created whole brain maps of the mean amplitude of low frequency fluctuations (mALFF) in middle-aged mouse lemurs. Then, we extracted mALFF in old lemurs to identify age-related changes. A high level of mALFF was detected in the temporal cortex (Brodmann area 20), somatosensory areas (Brodmann area 5), insula (Brodmann area 13-6) and the parietal cortex (Brodmann area 7) of healthy middle-aged mouse lemurs. Aging was associated with alteration of mALFF in somatosensory areas (area 5) and the parietal cortex (area 7).

**Graphical abstract:** 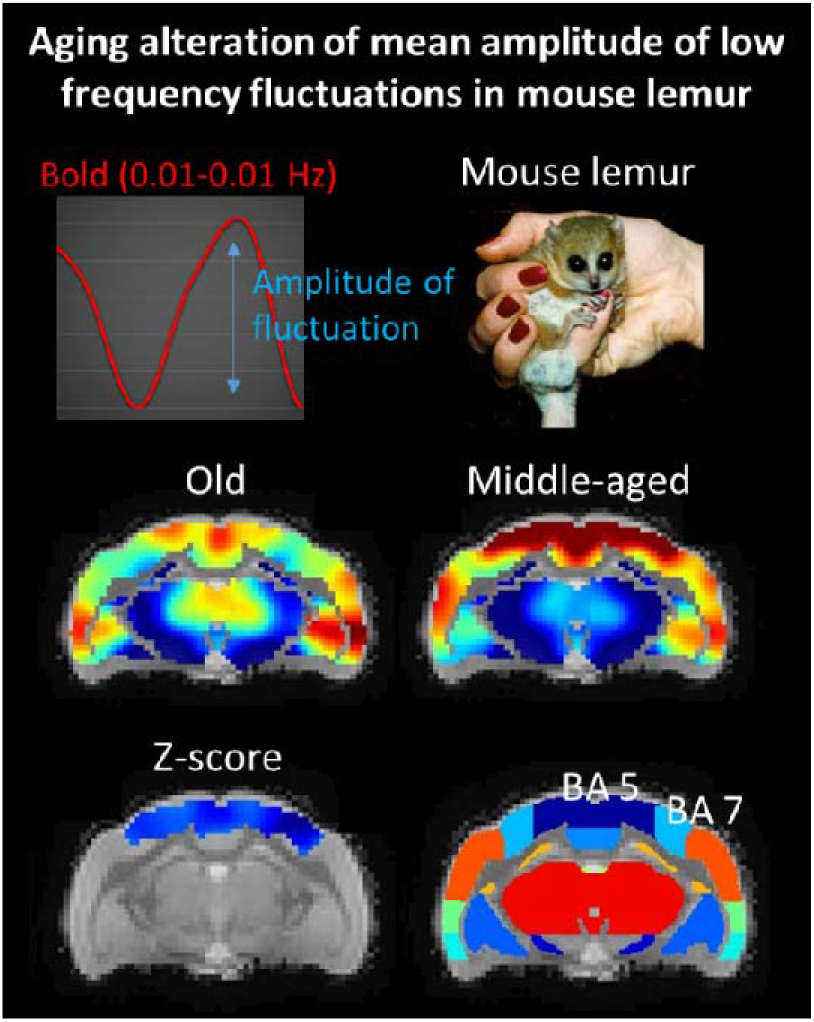

**Highlights:**

- We characterized mean amplitude of fluctuation at low frequencies (mALFF) in mouse lemurs.
- mALFF was the highest in regions involved in visuo-somatosensory-motor function (Brodmann areas 5, 7, 20) and in more integrative functions (area 13-16).
- mALFF was affected by aging in regions involved in visuo-somatosensory-motor function (parietal areas 5 and 7).
- mALFF is a useful marker to investigate age-related cerebral dysfunction in animals.

**Significance Statement:**

- The amplitude of low-frequency fluctuations (ALFF) is expected to reflect neuronal activity. It has been proposed as an MRI-based method to evaluate brain function.
- ALFF has been used to investigate different cerebral pathologies in animal models but the regional differences of ALFF signal and the impact of cerebral aging on ALFF has never been characterized.
- Here, we highlight for the first time regional difference of ALFF. High signal was detected in regions involved in visuo-somatosensory-motor function as well as in more integrative functions. ALFF was reduced in regions involved in visuo-somatosensory-motor function during aging. ALFF is thus a useful marker to investigate age-related cerebral dysfunction.

## 1. Introduction

Human life expectancy has dramatically increased during the last century. This comes with an increased risk for cerebral alterations leading to neurodegenerative diseases or mild cognitive/motor impairments that impair daily living. Monitoring functional impairments associated with cerebral ageing with translational imaging methods is critical to understand mechanisms leading to brain alterations and develop new treatments.

Measures of cerebral glucose metabolism using the radiolabeled glucose analog ^18^F-fluorodeoxyglucose (FDG) detected by Positron Emission Tomography (PET) imaging is a largely used marker to investigate brain function in elderlies (Edison et al., 2007; Kalpouzos et al., 2009) and in mouse model of Alzheimer’s disease (Franke et al., 2020). The use of PET is however restricted as this requires using radioactive compounds and is limited for animal studies by its low resolution when compared to functional magnetic resonance imaging (fMRI). In consequences, magnetic resonance imaging (MRI) is an interesting alternative to PET.

Low-frequency oscillations (LFO) of blood-oxygen level dependent (BOLD) signal reflects the total power of BOLD signal within the frequency range between 0.01 and 0.1 Hz. The amplitude of low-frequency fluctuations (ALFF) is expected to reflect neuronal activity (Zou et al., 2008) and was associated with markers of glucose metabolism (Aiello et al., 2015). Thus, it has been proposed as an MRI-based method to evaluate brain function (Biswal et al., 1995; Zou et al., 2008) and could be a promising radioactive-free alternative to FDG-PET. Studies in humans have shown that ALFF is negatively correlated with age in several brain regions such as the supplementary motor area, pre-supplementary motor area, anterior cingulate cortex, bilateral dorsal lateral prefrontal cortex, right inferior parietal lobule, and posterior cingulate cortex (Hu et al., 2014). However, healthy aging effects on ALFF indexes remain to our knowledge, unexplored in non-human mammalians.

The mouse lemur (*Microcebus murinus*) is a primate attracting increased attention in neuroscience research. This small animal (typical length 12cm, 60-120g weight) has a decade-long lifespan (Pifferi et al., 2018) and is a model for studying cerebral ageing (Sawiak et al., 2014) and Alzheimer’s disease (Kraska et al., 2011). This animal was used to establish the impact of prediabetes in the brain (Djelti et al., 2016) as well as to evaluate interventions modulating cerebral ageing process (Pifferi et al., 2018). The aim of the current study was thus to characterize ALFF in this primate in normal and ageing conditions. We described regional differences of ALFF signal and showed aged-related changes in specific brain regions.

## 2. Materials and methods

### 2.1. Animals and breeding

The guidelines of the European Communities Council directive (2010/63/EU) were followed when conducting this study. Our protocol was authorized by the local ethics committees CEtEA-CEA DSV IdF (authorizations 201506051736524 VI (APAFIS#778)). We originally included 33 mouse lemurs (21 males and 12 females) in our study. They were bred in our laboratory after being born at the CNRS/Brunoy, MNHN’s France, laboratory breeding colony (UMR 7179 CNRS/MNHN) (Molecular Imaging Research Center, CEA, Fontenay-aux-Roses). Four animals that displayed MR images with artefact or brain lesions were removed from the study.

<u>The “middle-aged lemur cohort”</u> consisted of 14 animals with an age range of 1.3 to 3.8 years (mean±SD: 2.1±0.8 years).

<u>The “old lemur cohort”</u> consisted of 15 animals with an age range of 8.0 to 10.8 years (mean±SD: 8.8±1.1 years).

The animals were housed in cages with one or two lemurs, enrichment for jumping and hiding, temperatures between 24 and 26 degrees Celsius, a relative humidity of 55 percent, and seasonal illumination (summer: 14 hours of light, 10 hours of darkness; winter: 10 hours of light, 14 hours of darkness). Food consisted of fresh apples and a handmade blend of bananas, cereals, eggs, and milk. Water supply for animals was freely accessible. None of the animals had ever taken part in invasive research or pharmaceutical trials before.

### 2.2. Animal preparation and MRI acquisition

To ensure animal stability during the experiment, all animals were scanned once while under isoflurane anesthesia at 1.25–1.5% in air, with respiratory rate monitoring. A 32°C air heating system was used to maintain body temperature, causing mouse lemurs to go into a state of natural torpor (Aujard, 2001). The benefit of this is that it prevents reawakening while maintaining a low anesthetic level. Animals were scanned on an 11.7 T Bruker BioSpec MRI machine (Bruker, Ettlingen, Germany) running ParaVision 6.0.1 with a volume coil for radiofrequency transmission and a quadrature surface coil for reception (Bruker, Ettlingen, Germany). We acquired anatomical images with a T2-weighted multi-slice multi-echo (MSME) sequence: TR = 5000 ms, TE = 17.5 ms, FOV = 32 × 32 mm, 75 slices of 0.2 mm thickness, 6 echoes, 5 ms IET, resolution = 200 μm isotropic, acquisition duration 10 min. We acquired resting state time series with a gradient-echo echo planar imaging (EPI) sequence: TR = 1000 ms, TE = 10.0 ms, flip angle = 90°, repetitions = 450, FOV = 30 × 20 mm, 23 slices of 0.9 mm thickness and 0.1 mm gap, resolution = 312.5 × 208.33 × 1000 μm, acquisition duration 7m30s.

### 2.3. MRI pre-processing

Data from scanners was exported as DICOM files and then changed to NIfTI-1 format. Then, using the Python program sammba-mri (SmAll MaMmals Brain MRI), spatial preprocessing was carried out (Celestine et al., 2020), http://sammba-mri.github.io) and we used nipype for pipelining (Gorgolewski et al., 2011), leverages AFNI (Cox, 1996) for most steps and RATS (Oguz et al., 2014) for “skullstripping”. A research template was made using the mutual registration of anatomical MR images. Images were then registered to a high-resolution anatomical mouse lemur template build for our previously published functional atlas (Garin et al., 2019). Motion, B0 distortion, and slice timing (interleaved) were removed from resting state MR images (per-slice registration to respective anatomical). Then, using sequential applications of the individual anatomical to study template and study template to mouse lemur atlas transformations, all the MR images were placed into the same space (the mouse lemur template). Functional images were further pre-treated using AFNI afni_proc.py (Cox, 1996). fMRI images were smoothed (0.9 mm), bandpass filtered, detrend corrected (0.01 to 0.1 Hz) as well as slice timing and motion corrected. TRs with excessive motion of 0.07 mm or where too many voxels were flagged as outliers by 3dToutcount (AFNI), were censored. To ensure steady-state magnetization, the first five volumes were not included in the study.

### 2.4. mALFF calculation and extraction

LFO measures were performed using the fast Fourier transform index: amplitude of low-frequency fluctuation (ALFF) (Zuo et al., 2010). As ALFF is sensitive to the scale of raw signal and the unit of BOLD signal is arbitrary, the original ALFF value is not adapted for comparisons between animals. In addition, ALFF can be susceptible to signal fluctuations caused by physiological noise unrelated to brain activity (Zou et al., 2008). A standardization procedure has been proposed by dividing the signal of each voxel by the global mean ALFF in each animal (Jia et al., 2020). The newly calculated index is called mean ALFF (mALFF). mALFF indexes were calculated for each voxel of the pre-processed EPI images in the low-frequencies range 0.01 to 0.1 Hz using the function “3dLombScargle” and “3dAmpToRSFC” from AFNI (Cox, 1996). The mALFF signal of each voxel was extracted within the different regions based on the anatomical atlas (Nadkarni et al., 2019) using NiftiLabelsMasker from Nilearn (Abraham et al., 2014).

### 2.5. Statistical analysis

Voxel wise analysis was performed using 3dttest++ from AFNI (Cox, 1996). This analysis identifies statistical differences between groups by using an uncorrected pvalue and a cluster size. Analysis space was reduced by thresholding the average mALFF map to the 20% highest voxels. Cluster size (856 voxels) was estimated on these remaining voxels using AFNI (-Clusterize) with p<0.05 (uncorrected). Then, using “map_threshold” from nilearn (Abraham et al., 2014), we extracted on the Ttest map any cluster of statistical voxels superior to 856 associated to p<0.05.

## 3. Results

### 3.1. mALFF in middle-aged mouse lemurs

mALFF maps were recorded from 14 middle-aged mouse lemurs. Individual mALFF maps of each animal were averaged to produce 3D maps of the group (Fig. 1). Automatic extractions of the mALFF signal were then performed in various cerebral regions by using a reference anatomical atlas ((Nadkarni et al., 2019); Fig. 1A). The highest cortical signal was observed in the temporal cortex (Brodmann area (BA) 20 *i.e*. secondary visual area expected to be involved in visual processing and recognition memory), parietal cortex (BA 5 *i.e*. a secondary somatosensory areas; BA 7 *i.e*. an integrative area involved in visuo-motor coordination). High signal was also detected in the primary somatosensory cortex (BA 1-3) and the primary motor cortex (BA 4). More integrative areas as the insula (BA 13-16) and in the cingulate cortex (BA 23 and 24) also displayed high signal. In subcortical areas, the basal forebrain exhibited the most elevated signal when compared to the rest of the brain. Conversely, cortical regions such as BA 30 (agranular retrolimbic area involved in vision) and the adjacent BA 27 (area presubicularis involved in vision) or area 25 (antero-ventral part of the cingulate cortex) displayed the lowest levels of mALFF. Subcortical regions such as substantia nigra, subthalamic nucleus or hypophysis also displayed the lowest levels of mALFF.

**Figure 1.**
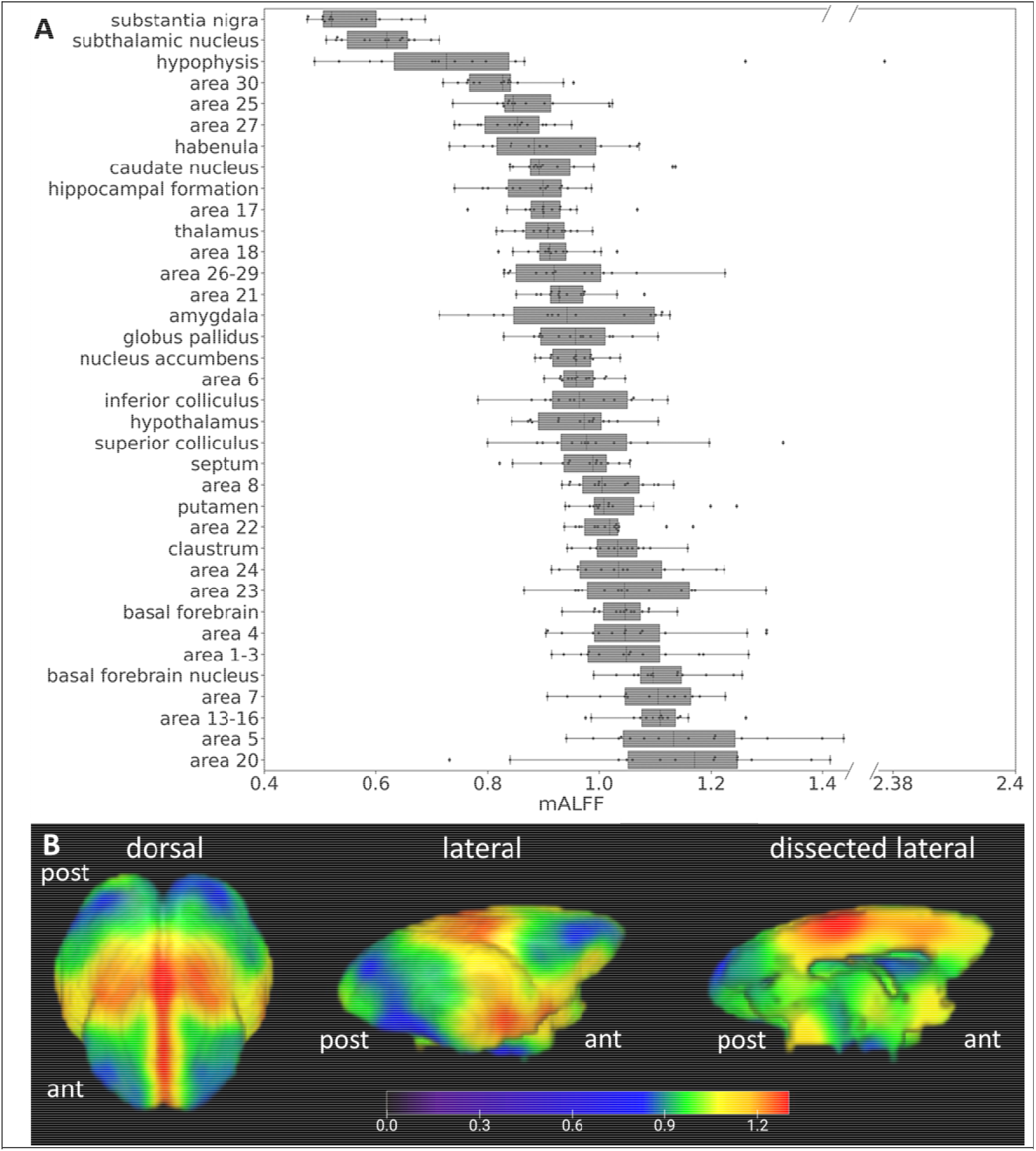
Whole brain mALFF average-map in middle-aged mouse lemurs. (A) mALFF in different brain regions of middle-aged adult mouse lemurs (n=14) ranked based on their group median value. Elevated mALFF is observed within regions encompassing cortical Brodmann areas 20, 5, 13-16, 7, 1-3, 4, 23, 24 and basal forebrain. 3D surface average-map of the mALFF showing regional signal variations (B). ant: anterior part of the brain, post: posterior part of the brain.

### 3.2. Age-related changes of mALFF

BOLD signal was recorded from 15 old mouse lemurs (mean age±SD: 8.8±1.1 years) and as for middle-aged lemurs, mALFF analysis was performed. Average mALFF maps are displayed in Fig. 2A for middle-aged mouse lemurs and in Fig. 2B for old mouse lemurs. Comparison of average mALFF map between middle-aged and aged lemurs indicates lower mALFF in the parietal cortex of aged animals and a trend toward higher signal in their hypothalamic regions. Voxelwise analysis between groups revealed a significant loss of BOLD signal amplitude in parietal regions involving (BA 5 and BA 7) of aged animals (Fig 2C, D).

**Figure 2.**
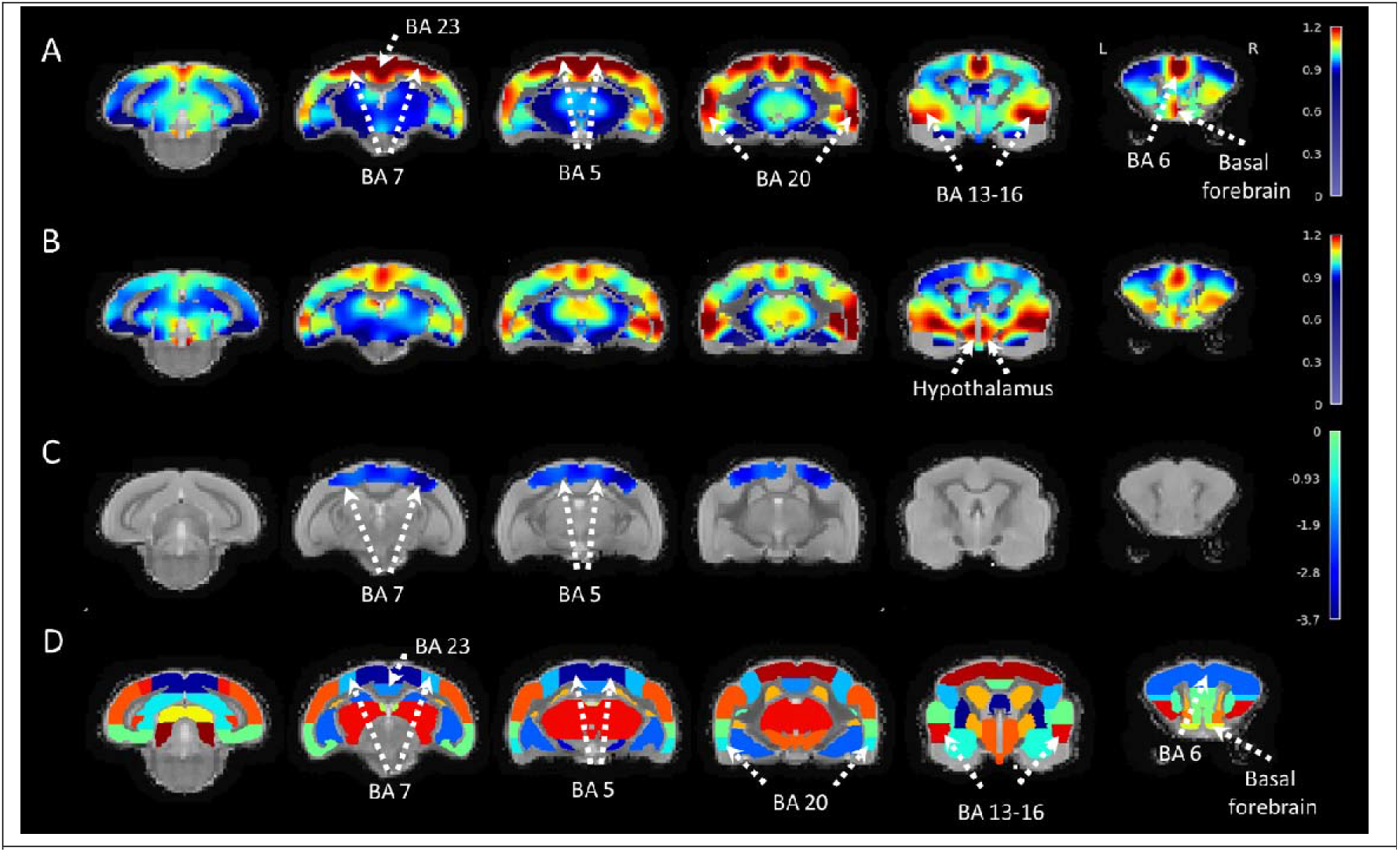
mALFF contrast in middle-aged and old mouse lemurs. Average mALFF in middle-aged (A) and old mouse lemurs (B). Significant statistical decrease of the mALFF signal was detected using voxelwise analysis in the Brodmann areas 5 and 7 (C). Anatomical view of the mouse lemur atlas corresponding to the displayed sections (A-C) are shown in D (Nadkarni et al., 2019). BA: Brodmann area. Scale bars in A-C: mALFF contrast, in D: z-score.

## 4. Discussion

### 4.1. Cerebral distribution of mALFF index in mouse lemurs

This study evaluated mALFF index in mouse lemur primates at high field (11.7T). The evaluation of mALFF in a middle-aged cohort provided 3D maps of the normal distribution of the mALFF indexes. Highest levels of mALFF were detected in cortical structures involved in secondary visual (area 20 in temporal region), somatosensory (area 5 in parietal region), and in motor processing and integrative regions involved in visuo-motor coordination (area 7 in parietal region). High signal was also detected in the primary somatosensory cortex (BA 1-3) and the primary motor cortex (BA 4). More integrative areas as the insula (BA 13-16) and in the cingulate cortex (BA 23 and 24) also displayed high signal.

### 4.2. Age-related changes of mALFF index

As a second part of the study, we assessed age-related changes of mALFF. In humans, healthy ageing mainly affects ALFF in brain regions involved in motor function such as the supplementary motor area (BA 5) or the pre-supplementary motor area. More integrative regions such as the anterior cingulate cortex, the dorso-lateral prefrontal cortex, the posterior cingulate cortex, and the inferior parietal lobule are also impaired (Hu et al., 2014). In mouse lemurs, the only regions significantly affected by healthy ageing were those involved in visuo-somatosensory-motor function (parietal regions as BA 5 *i.e*. a secondary somatosensory areas and BA 7 *i.e*. an integrative area involved in visuo-motor coordination). These two regions are involved in somatosensory processing, in movement such as grasping (Gardner et al., 2007) or object location (Caminiti et al., 2010). These alterations could participate to the described visuo-somatosensory-motor alterations reported in aged lemurs (Le Brazidec et al., 2017).

Atrophy detected in mouse lemurs impacts the whole brain in the latest stages of ageing but the most prominent atrophied areas include the insular (areas 13-16), frontal (area 6), parietal (areas 5, 7), occipital (areas 17, 18), inferior temporal (areas 21, 28) and cingulate cortices (areas 23, 24, 25) (Nadkarni et al., 2019; Sawiak et al., 2014). In consequence, an ageing effect can be detected with mALFF and anatomical measures in both areas 5 and 7 but areas such as 17, 18, 13-16 are impacted only by atrophy and not by mALFF changes. A better understanding of the significance of mALFF changes is now needed to further interpret the significance of mALFF changes.

In previous studies, ALFF or fractional ALFF were used to evaluate the effect of simian immunodeficiency virus (Zhao et al., 2019), spinal injury (Rao et al., 2014) or anesthesia (Wu et al., 2016) in macaques. In mice, Huntington-related pathological effects were detected using ALFF (Chang et al., 2018). In rats, various effects of stress could also be evaluated using ALFF (Li et al., 2018; Yan et al., 2017). By showing age-related changes, the current study increases the range of ALFF changes detected in mammals in pathological situations.

### 4.3. Limitations

Anesthesia is known to interact with brain function. For example, Isoflurane changes functional connections between brain regions in marmosets (Garin et al., 2022). Thus, one of the possible limitations of the study is anesthesia that could modulate mALFF signal and lead to whole brain mALFF average-map that does not reflect mALFF in non-anesthetized animals. Future studies will thus have to be conducted to investigate the impact of anesthesia on mALFF.

Comparison between middle-aged and old animals revealed age-related differences of mALFF. As both groups were anesthetized, this difference is expected to reflect the ageing effect. We can however not exclude that anesthesia may have decreased our ability to detect mALFF changes in some regions affected by ageing.

## 5. Conclusion

In conclusion, this study provides the proof of concept that mALFF can be measured in the whole brain of mouse lemurs and can detect aged-related changes. It proposes mALFF as a tool for the exploration of the cerebral function in mammals as well as an interesting candidate for the longitudinal follow up of age-related cerebral dysfunction animal models.

## 6. Acknowledgements

We thank the France-Alzheimer Association, Plan Alzheimer Foundation and the French Public Investment Bank’s “ROMANE” program for funding this study. The 11.7T MRI scanner was funded by a grant from NeurATRIS: A Translational Research Infrastructure for Biotherapies in Neurosciences (“Investissements d’Avenir”, ANR-11-INBS-0011). C.G. was financed by the French Ministère de l’Enseignement Supérieur, de la Recherche et de l’Innovation.

## 7. Competing interests

The authors do not have financial and non-financial competing interests in relation to the work described.

## 8. Author contributions

C.M.G. and M.D. contributed to the study conception and design. C.M.G. designed the BOLD sequences, C.M.G. designed registrations strategies and pipelines. C.M.G. and M.D. wrote the manuscript.

## 9. Data and code availability statements

### Availability of the data used in the study

Raw MRI data in mouse lemurs are available upon request following a formal data sharing agreement required by authors’ institution.

The template and atlas used in this study maps are available for download in NIfTI-1 format at https://www.nitrc.org/projects/mouselemuratlas.

### Availability of the software data used in the study

All the software used to perform the analysis are publicly available. In particular, sammba-mri, the code developed by our team for spatial pre-processing is available at http://sammba-mri.github.io. All the third party codes used in the article are cited.

## Notes

### Competing Interest Statement

The authors have declared no competing interest.

### Summary of Updates

update of the material and method

